# Critical-Period Visual Deprivation Disrupts Binocular Integration but Spares Spatial Acuity in the Geniculocortical Pathway

**DOI:** 10.1101/484774

**Authors:** Carey Y. L. Huh, Karim Abdelaal, Kirstie J. Salinas, Diyue Gu, Jack Zeitoun, Dario X. Figueroa Velez, John P. Peach, Charless C. Fowlkes, Sunil P. Gandhi

**Affiliations:** Department of Neurobiology and Behavior, University of California, Irvine, California 92697; School of Biological Sciences, University of California, Irvine, California 92697; Donald Bren School of Information & Computer Sciences, University of California, Irvine, California 92697; Whiting School of Engineering, Johns Hopkins University, Baltimore, Maryland 21218; Department of Computer Science, University of California, Irvine, California 92697; Center for Neurobiology of Learning and Memory, University of California, Irvine, California 92697

**Keywords:** Thalamus, Dorsolateral Geniculate Nucleus, Visual Cortex, Critical Period, Binocular Vision, Spatial Acuity, GCaMP6, Amblyopia

## Abstract

Monocular deprivation (MD) during the juvenile critical period leads to long-lasting impairments in binocular function and visual acuity. The site of these changes has been widely considered to be cortical. However, recent evidence indicates that binocular integration may first occur in the dorsolateral geniculate nucleus of the thalamus (dLGN), raising the question of whether MD during the critical period may produce long-lasting deficits in dLGN binocular integration. Using *in vivo* two-photon Ca^2+^ imaging of dLGN afferents and excitatory neurons in superficial layers of primary visual cortex (V1), we demonstrate that critical-period MD leads to a persistent and selective loss of binocular dLGN inputs, while leaving spatial acuity in the thalamocortical pathway intact. Despite being few in number, binocular dLGN boutons display remarkably robust visual responses, on average twice stronger than monocular boutons, and their responses are exquisitely well-matched between the eyes. To our surprise, we found that MD leads to a profound binocular mismatch of response amplitude, spatial frequency and orientation tuning detected at the level of single thalamocortical synapses. In comparison, V1 neurons display deficits in both binocular integration and spatial acuity following MD. Our data provide the most compelling evidence to date demonstrating that following critical-period MD, binocular deficits observed at the level of V1 may at least in part originate from dLGN binocular dysfunction, while spatial acuity deficits arise from cortical circuits. These findings highlight a hitherto unknown role of the thalamus as a site for developmental refinement of binocular vision.

---

SIGNIFICANCE STATEMENT

Abnormal binocular vision is a hallmark of amblyopia, a disorder that affects 2 – 5% of the population. Recent evidence suggests that information from the two eyes combines at an earlier stage than the visual cortex – in the thalamus. It is unknown whether binocular integration in the thalamus can be permanently altered by manipulations of early visual experience. Using *in vivo* two-photon calcium imaging, we show that depriving one eye of input during a sensitive period in development chronically and profoundly impairs binocular integration in the thalamus. This discovery sheds new light on the potential role for developmental mechanisms in the thalamus in establishing binocular vision.

## INTRODUCTION

Long-term monocular deprivation (MD) during the critical period of visual system development chronically alters ocular dominance (1–3), impairs spatial acuity (4–6) and disrupts binocular matching of visual properties (7–9). Abnormal binocular vision and reduced acuity are the hallmarks of amblyopia, a visual disorder that arises from unbalanced binocular input during early childhood (10). Early work using MD models demonstrated profound functional changes in primary visual cortex (V1) that were not detected in the dorsolateral geniculate nucleus of the thalamus (dLGN) (11–13). However, thalamocortical projections have been observed to undergo anatomical changes in MD models (14–17), and recent brain imaging studies indicate that human amblyopes display anatomical and functional thalamic deficits (18, 19). Given these lines of evidence, the impact of critical-period MD on functional properties of dLGN requires re-examination.

A growing body of recent evidence indicates that significant binocular processing occurs in dLGN in mice (20–22) and marmosets (23). A retrograde tracing study (24) demonstrated that single dLGN neurons receive direct retinal inputs from both eyes, providing an anatomical substrate for binocular integration in the mouse dLGN. A functional study (21) showed that ocular dominance of thalamic axons in V1 can be altered by short-term (6-8 days) MD under plasticity-enhancing conditions in adult mice but the effect was found to be transient. Another study (22) reported that seven days of MD during the critical period can lead to an ocular dominance shift in dLGN neurons. However, these studies did not address whether long-term MD during the critical period, a manipulation that has been shown to have a long-lasting impact on the animal’s visual acuity (25–27), can lead to persistent changes in dLGN visual properties, including binocular integration.

Using *in vivo* two-photon Ca^2+^ imaging, we investigated the visual responses of dLGN axons in V1 in adult mice that underwent long-term (14-day) MD during the critical period *vs.* control littermates. We found that MD leads to a persistent loss of binocularity and interocular mismatch of visual properties in dLGN inputs while leaving spatial acuity intact in the thalamocortical pathway. In contrast, MD led to reductions in both binocularity and visual acuity in V1 L2/3 excitatory neurons. These findings indicate that binocularity deficits associated with amblyopia may originate from thalamic binocular dysfunction, while spatial acuity deficits may be cortically based.

## RESULTS

### Selective loss of binocular thalamocortical boutons following critical-period MD

In order to target the expression of the calcium sensor GCaMP6s to thalamocortical projections from relay neurons in dLGN, we injected a Cre-dependent GCaMP6s virus into dLGN in VGLUT2-Cre mice (Fig. 1A; see Materials and Methods). We performed calcium imaging in adult mice (P93 – 132, mean: P106) that were either monocularly deprived for 14 days during the critical period (P19-33) or littermate controls (Fig. 1B). Calcium imaging was performed in awake mice that were viewing drifting gratings of various orientations and spatial frequencies (Fig. 1C). Two-photon Ca^2+^ imaging was performed in superficial layers (L1-2/3) of bV1 (Fig. 1D-G). In deprived mice, dLGN injections and functional imaging were performed in the hemisphere contralateral to the deprived eye.

**Figure 1.**
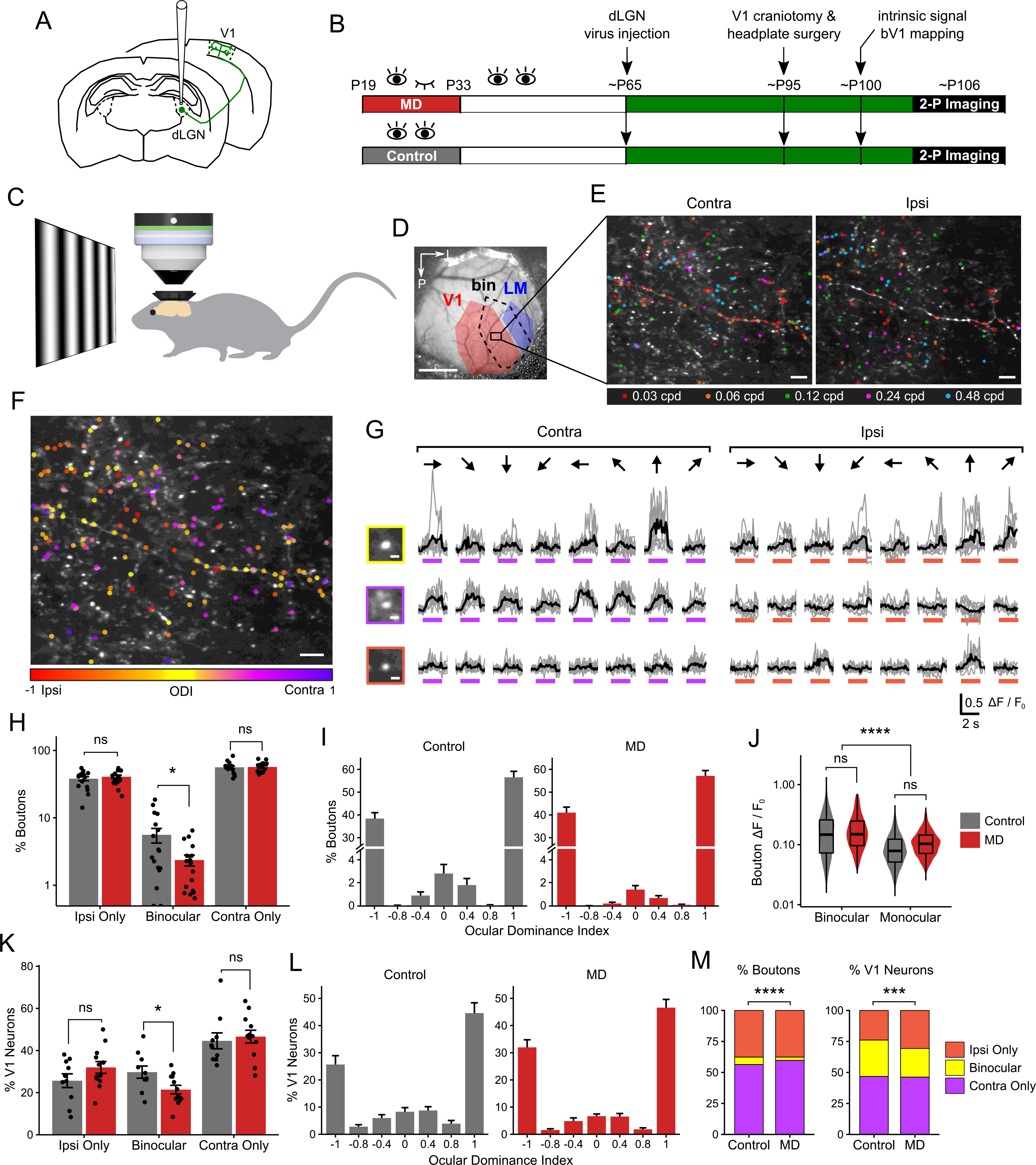
Selective loss of binocular thalamocortical boutons following critical-period MD. **A**, Schematic of dLGN virus injection and GCaMP6s expression in thalamocortical axons in V1. **B**, Experimental timeline. **C**, *In vivo* two-photon Ca^2+^ imaging of visual responses were performed in awake head-fixed mice. **D**, An example cranial window with the binocular zone mapped using widefield intrinsic signal imaging (scale bar 1 mm). **E**, An example field of view (summed projection) of dLGN boutons imaged in bV1 L1-2/3 of a control mouse. Visually responsive boutons (see Materials and Methods for criteria) were color-coded according to peak SF during contralateral-(left) and ipsilateral-eye (right) viewing (scale bar 10 μm). m). **F**, Same dataset as in **E** but color-coded for ocular dominance (see Materials and Methods). **G**, Ca^2+^ signals in a binocular (top) and two monocular (middle, bottom) example boutons in response to drifting gratings presented to contralateral or ipsilateral eye (gray: individual traces, black: mean trace, purple and orange bars: time of stimulus presentation; bouton image scale bar 2 μm). m). Responses to 8 orientations at peak SF are shown. **H**, Percentage of visually responsive dLGN boutons per field in control *vs.* MD mice (mean ± SEM per field, *n* = 17 control, 17 MD fields; linear mixed-effects model, effect of MD for ipsi-only: *P* = 0.67, binocular: *P* = 0.04, contra-only: *P* = 0.90). **I**, Ocular dominance index distribution of boutons in control *vs.* MD mice (mean ± SEM per field, *n* = 17 control, 17 MD fields). **J**, Violin and overlaid box plots of mean response amplitudes (R_pref_) of boutons (linear mixed-effects model, effect of MD: *P* = 0.50, binocular *vs.* monocular: *P* = 2.2 x 10^−16^). In box plots, the central mark indicates the median and the bottom and top edges indicate the 25^th^ and 75^th^ percentiles, respectively. **K**, Percentage of visually responsive V1 L2/3 excitatory neurons per field in control *vs.* MD mice (mean ± SEM per field, *n* = 10 control, 12 MD fields; linear mixed-effects model, effect of MD for ipsi-only: *P* = 0.14, binocular: *P* = 0.01, contra-only: *P* = 0.66). **L**, Ocular dominance index distribution of V1 neurons in control and MD mice (mean ± SEM per field). **M**, Fraction of visually responsive dLGN boutons (left) and V1 neurons (right) that are ipsi-only, contra-only or binocular (boutons: χ^2^(2) = 38.3, *P* = 4.8 x 10^−9^, *n* = 2866 boutons in 5 control mice, 2975 boutons in 6 MD mice; V1 neurons: χ^2^(2) = 17.4, *P* = 1.6 x 10^−3^, *n* = 1051 neurons in 9 control mice, 1145 neurons in 4 MD mice). All panels: ^ns^ *P* > 0.05, \**P* < 0.05, \*\*\**P* < 0.001, \*\*\*\**P* < 0.0001.

We found that most dLGN boutons in superficial layers of bV1 were monocular (visually responsive to contralateral or ipsilateral eye only) with a small fraction of boutons displaying significant visual responses to both eyes (binocular; 6% in control mice; Fig. 1E-J, 1M; see also Supplemental Movie 1). To our surprise, we found that critical-period MD led to a long-lasting reduction in the number and relative proportion of binocular dLGN boutons recorded per field of view (Fig. 1H-I and Fig. S1A). This was not due to reduced detectability of binocular boutons in MD mice, because there was no significant difference in response amplitudes of boutons between control and MD mice (Fig. 1J and Fig. S1B). Interestingly, binocular boutons displayed approximately twice greater response amplitudes compared to monocular boutons (median R_pref_ for binocular boutons: 0.15, monocular boutons: 0.08). MD significantly reduced the binocular fraction among dLGN boutons from 6% to 3% in control *vs.* MD mice (Fig. 1M, left; see also Fig. S1F-G). The number of virally infected neurons and their spatial distribution in dLGN were comparable between functionally imaged control and MD mice and could not account for the reduction in binocular boutons (Fig. S2).

For comparison, we also examined the impact of critical-period MD on L2/3 excitatory neurons in bV1 by performing two-photon Ca^2+^ imaging from GCaMP6s-expressing cells in CaMK2a-tTA;tetO-GCaMP6s transgenic mice (28). We found that MD led to a shift in ocular dominance distribution (Fig. S1C) and a reduction in the percentage of binocular bV1 neurons from 29% to 23% in control *vs.* MD mice (Fig. 1K-M) without a significant reduction in response amplitude (Fig. S1D-E). These results suggest that MD-induced binocularity deficits observed at the level of V1 may originate, at least in part, from binocular dLGN input loss.

### Intact SF processing in thalamocortical boutons following critical-period MD

Following critical-period MD, mice develop reduced visual acuity in the deprived eye, an impairment that lasts well into adulthood (25–27). It remains unclear whether acuity deficits are generated *de novo* in cortical circuits or are inherited from dLGN. It is also unknown how spatial frequency (SF) representation interacts with ocular dominance in dLGN. Previously, we showed that contralateral-eye dominated monocular V1 neurons prefer higher SF compared to binocular neurons (29). Thus, we explored how binocularity and SF processing interact in dLGN boutons and whether critical-period MD affects these properties.

We found that dLGN boutons were tuned to a wide range of SF from 0.03 to 0.96 cpd (Fig. 2A-D and Fig. S3-4). Many dLGN boutons exhibited sharp SF tuning (Fig. 2B-C). As a population, dLGN boutons were tuned to higher SF compared to V1 neurons (Fig. 2D-H and Fig. S4A-B), consistent with a previous report using electrophysiological recordings (30). Binocular boutons were tuned to significantly lower SF compared to contralateral-eye dominated monocular boutons (Fig. S4A-B), similar to our observations in V1 neurons.

**Figure 2.**
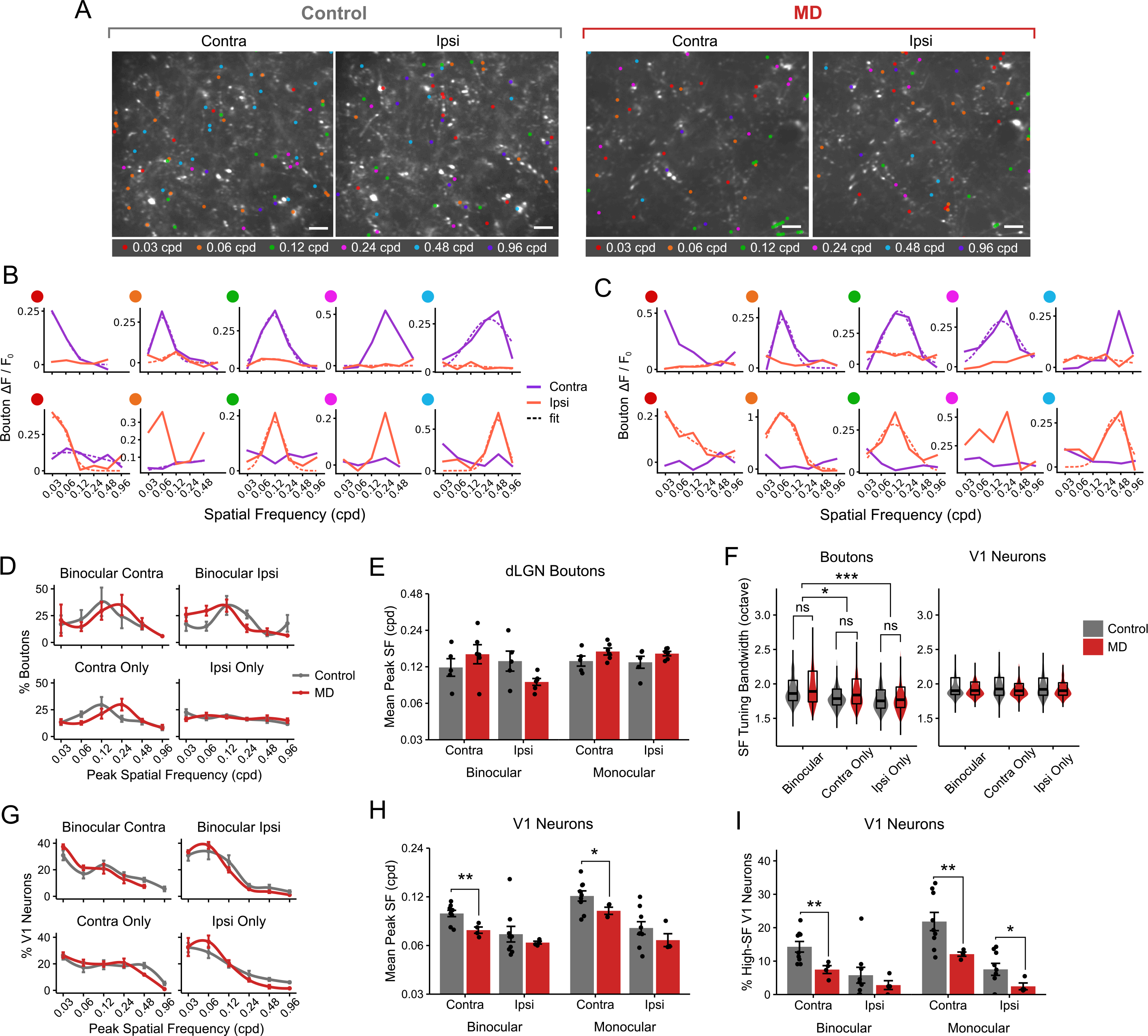
Intact SF processing in thalamocortical boutons following critical-period MD. **A**, Example fields of view of dLGN boutons imaged in bV1, color-coded according to peak SF of bouton during contralateral-*vs.* ipsilateral-eye presentation in control and MD mice (scale bar 10 μm). m). **B**, Example SF tuning curves of boutons that were responsive to contralateral eye (top) and ipsilateral eye (bottom) in control mice. Purple: contralateral-eye trials; orange: ipsilateral-eye trials; solid: mean response amplitudes; dotted: fitted curves based on mean values. Fits are omitted if curve-fitting failed to converge. Only monocular boutons are shown. **C**, Example SF tuning curves of boutons in MD mice. **D**, Mean probability distribution of peak SF in dLGN boutons, shown separately for binocular-contra, binocular-ipsi, monocular-contra, monocular-ipsi responses (mean ± SEM of by-animal values, *n* = 5 control *vs.* 6 MD mice). **E**, Mean peak SF of boutons in control *vs.* MD mice (mean ± SEM of by-animal values, 3-way ANOVA, control *vs.* MD: *P* = 0.56, binocular *vs.* monocular: *P* = 0.04, contra *vs.* ipsi: *P* = 0.13). **F**, Violin and overlaid box plots of SF bandwidth in binocular, contra-only *vs.* ipsi-only monocular dLGN boutons (left) and V1 neurons (right) in control and MD mice. Linear mixed-effects model. Boutons: effect of MD: *P* = 0.08, binocular *vs.* contra-only: *P* = 0.01, binocular *vs.* ipsi-only: *P* = 0.0009, contra-only *vs.* ipsi-only: *P* = 0.15. V1 neurons: effect of MD: *P* = 0.54, binocular *vs.* contra-only: *P* = 0.68, binocular *vs.* ipsi-only: *P* = 0.68, contra-only *vs.* ipsi-only: *P* = 0.59. In box plots, the central mark indicates the median and the bottom and top edges indicate the 25^th^ and 75^th^ percentiles, respectively. **G**, Mean probability distribution of peak SF in V1 L2/3 excitatory neurons (mean ± SEM of by-animal values, *n* = 9 control *vs.* 4 MD mice). Note leftward shift of SF distribution curves in MD mice compared to controls. For **D** and **G**, mean values were fitted with a local regression smoothing function. **H**, Mean peak SF of V1 L2/3 neurons in control *vs.* MD mice (mean ± SEM of by-animal values, 3-way ANOVA, effect of MD: *P* = 0.007, binocular *vs.* monocular: *P* = 0.01, contra *vs.* ipsi: *P* < 10^−6^; post-hoc tests: effect of MD in binocular-contra: *P* = 0.004, binocular-ipsi: *P* = 0.32, monocular-contra: *P* = 0.03, monocular-ipsi: *P* = 0.21). **I**, Percentage of V1 neurons with peak SF of 0.48 – 0.96 cpd (mean ± SEM of by-animal values, 3-way ANOVA, effect of MD: *P* = 0.0007, binocular *vs.* monocular: *P* = 0.01, contra *vs.* ipsi: *P* = 7.6 x 10^−8^; post-hoc tests: effect of MD in binocular-contra: *P* = 0.005, binocular-ipsi: *P* = 0.30, monocular-contra: *P* = 0.007, monocular-ipsi: *P* = 0.03. All panels: ^ns^ *P* > 0.05, \**P* < 0.05, \*\**P* < 0.01, \*\*\**P* < 0.001.

Interestingly, we found no statistically significant effect of critical-period MD on dLGN boutons’ preferred SF (Fig. 2D-E). SF tuning bandwidths of dLGN boutons were comparable between control and MD mice and similar to those found in V1 neurons (Fig. 2F). In V1 neurons, however, MD led to a robust reduction in preferred SF, particularly for contralateral (deprived) eye responses (Fig. 2G-H). Significantly fewer V1 neurons preferred 0.48 – 0.96 cpd in MD mice compared to controls (Fig. 2G, 2I). These results indicate that critical-period MD impairs visual acuity in V1 neurons, while leaving SF processing in dLGN responses intact. It suggests that binocular vision during the critical period is necessary for the development of visual acuity in V1 but not in dLGN.

### Binocular mismatch in thalamocortical boutons following critical-period MD

Critical-period MD has been shown to lead to long-lasting binocular mismatch of orientation tuning in V1 neurons (8, 31) but whether the mismatch originates from dLGN inputs has been unclear. In addition, it is unknown whether the binocular mismatch extends to other visual properties such as SF tuning. Thus, we investigated whether MD leads to interocular mismatch in binocular dLGN responses, in terms of preferred SF and orientation, at the level of single thalamocortical synapses in V1.

In controls, binocular dLGN boutons were exquisitely well-matched between the eyes in terms of response amplitude and preferred SF (Fig. 3A-G). The majority (54%) of binocular boutons displayed ocular dominance index (ODI) values between −0.2 and +0.2 (Fig. 3C) and 40% of binocular boutons showed exact peak SF matching between the eyes (Fig. 3E-G). Among SF-mismatched boutons, preferred SF was higher in contralateral- or ipsilateral-eye responses in approximately equal proportions of boutons (Fig. 3E, 3G). Binocular boutons also exhibited significant matching in terms of preferred orientation (Fig. 3H-J).

**Figure 3.**
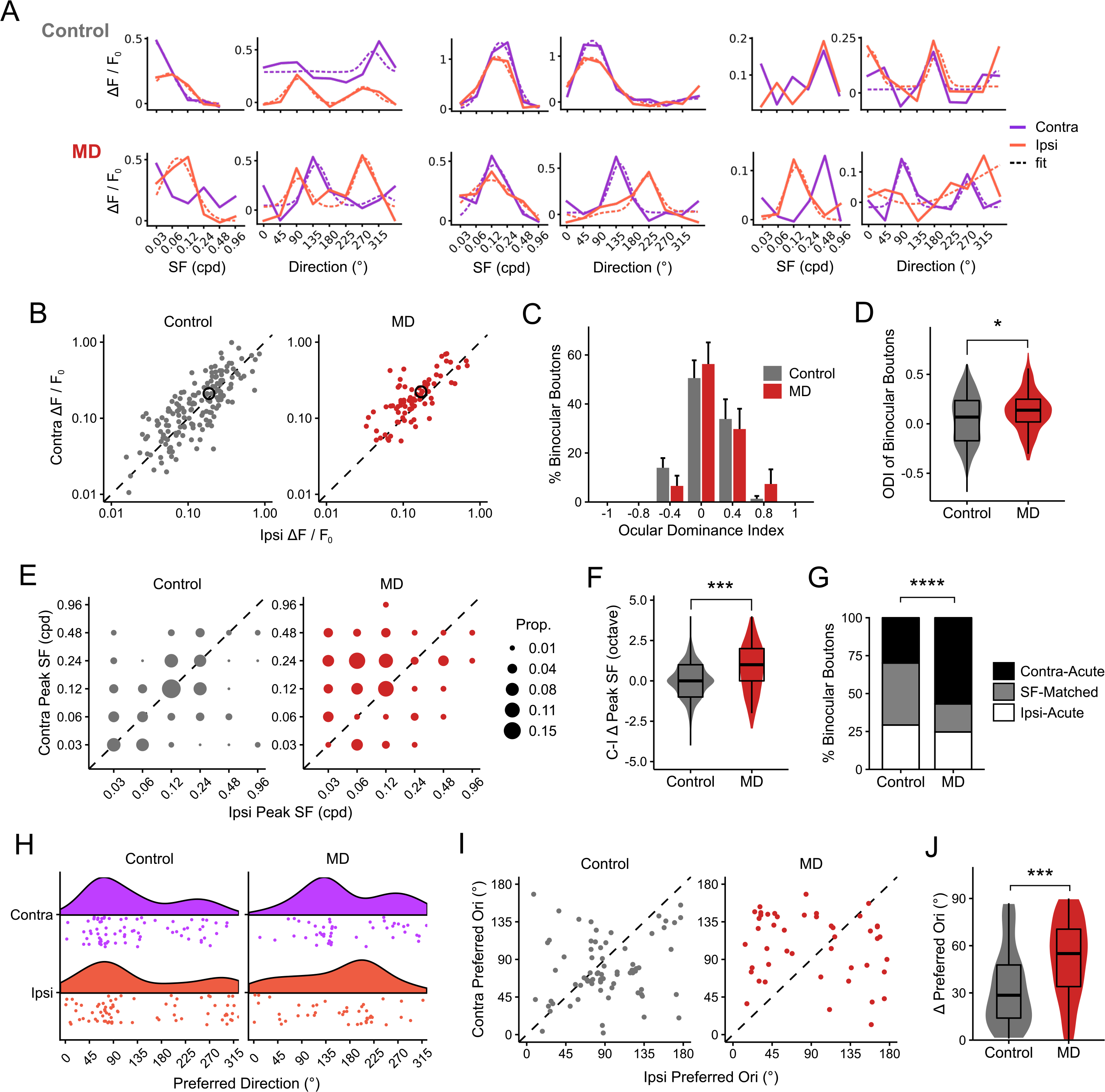
Binocular mismatch in thalamocortical boutons following critical-period MD. **A**, Each pair of plots show SF tuning curve (left) and orientation tuning curve at peak SF (right) of the same binocular bouton to illustrate example tuning curves in control (top 3 boutons) and MD (bottom 3 boutons) mice. Purple: contralateral-eye trials; orange: ipsilateral-eye trials; solid: mean response amplitudes; dotted: fitted curves based on mean values. Fits are omitted if curve-fitting failed to converge. Only binocular boutons are shown. **B**, Visual response amplitudes of binocular dLGN boutons to preferred SF and orientation (R_pref_) during contralateral-(y-axis) *vs.* ipsilateral-eye (x-axis) viewing. Black open circles indicate mean values. **C**, Ocular dominance distribution of binocular boutons (mean ± SEM per field, *n* = 17 fields in 5 control mice, 17 fields in 6 MD mice). **D**, Ocular dominance index values of binocular boutons in control *vs.* MD mice (Wilcoxon rank sum test: *P* = 0.02). **E**, Proportion plots of contralateral-*vs.* ipsilateral-eye peak SF in binocular boutons in control *vs.* MD mice. Unity (dotted line) represents perfect match. **F**, Interocular difference in peak SF (contra – ipsi) for binocular boutons in control *vs.* MD mice (Wilcoxon rank sum test: *P* = 0.0008). **G**, Fractions of binocular boutons that were SF-Matched, Contra-Acute (peak SF is greater in contralateral-eye response) or Ipsi-Acute (peak SF is greater in ipsilateral-eye response) in control *vs.* MD mice (Chi-squared test: χ^2^(2) = 18.9, *P* = 7.6 × 10^−5^). **H**, Rain cloud plots showing distributions of preferred direction in orientation-or direction-selective (OS/DS; gOSI or gDSI > 0.25) binocular boutons in control *vs.* MD mice. **I**, Scatter plot of preferred orientation of binocular boutons during contralateral-*vs.* ipsilateral-eye viewing. **J**, Interocular difference in preferred orientation for binocular boutons in control *vs.* MD mice (Wilcoxon rank sum test: *P* = 0.00027). **B-G**: *n* = 171 control *vs.* 81 MD binocular boutons. **H-J**: *n* = 74 control *vs.* 46 MD OS/DS binocular boutons. For all panels: *n* = 5 control vs. 6 MD mice, \**P* < 0.05, \*\*\**P* < 0.001.

We found that critical-period MD led to a greater binocular mismatch in visual properties. In MD mice, binocular boutons were less well-matched in response amplitude between the eyes and this was reflected in a small but significant change in ODI of binocular boutons towards the contralateral (deprived) eye (ODI shift: +0.09; Fig. 3C-D). We also found a marked binocular mismatch in preferred SF in MD mice, with ipsilateral (non-deprived) eye responses being tuned to lower SF compared to contralateral-eye responses in MD mice (Fig. 3E-G). The SF mismatch was not due to boutons exhibiting broader SF tuning in MD mice (Fig. 2F). MD also led to a higher degree of binocular mismatch in preferred orientation in orientation- or direction-tuned binocular dLGN boutons (Fig. 3H-J). Overall, many dLGN boutons were highly direction-tuned (Fig. 3A and Fig. S3, S4C-D), consistent with previously reported properties of dLGN neurons projecting to superficial layers of V1 (32–36). Orientation/direction selectivity indices were similar between control and MD mice (Fig. S4E-F). These findings indicate that normal binocular vision during the critical period of development is necessary for proper binocular matching of visual properties, including response amplitude, SF and orientation tuning, in dLGN neurons.

### No gross structural loss of thalamocortical connectivity following critical-period MD

Considering that binocular dLGN inputs constitute a relatively small proportion of the total thalamocortical input and they are selectively impaired following critical-period MD (Fig. 1), we hypothesized that there would be little to no gross structural deficit in thalamocortical projections following MD. Indeed, we found no significant long-lasting alterations in the overall density and thickness of dLGN axons in V1 L1-2/3 following critical-period MD (Fig. 4), consistent with previous studies that used 7- or 20-day MD and found no chronic morphological changes (16, 17). This suggests that critical-period visual experience is essential for the development of normal visual function, rather than structure, of overall dLGN projections to superficial layers of V1. However, future studies are needed to dissect whether the form, as well as function, of binocular thalamocortical projections are impaired by MD.

**Figure 4.**
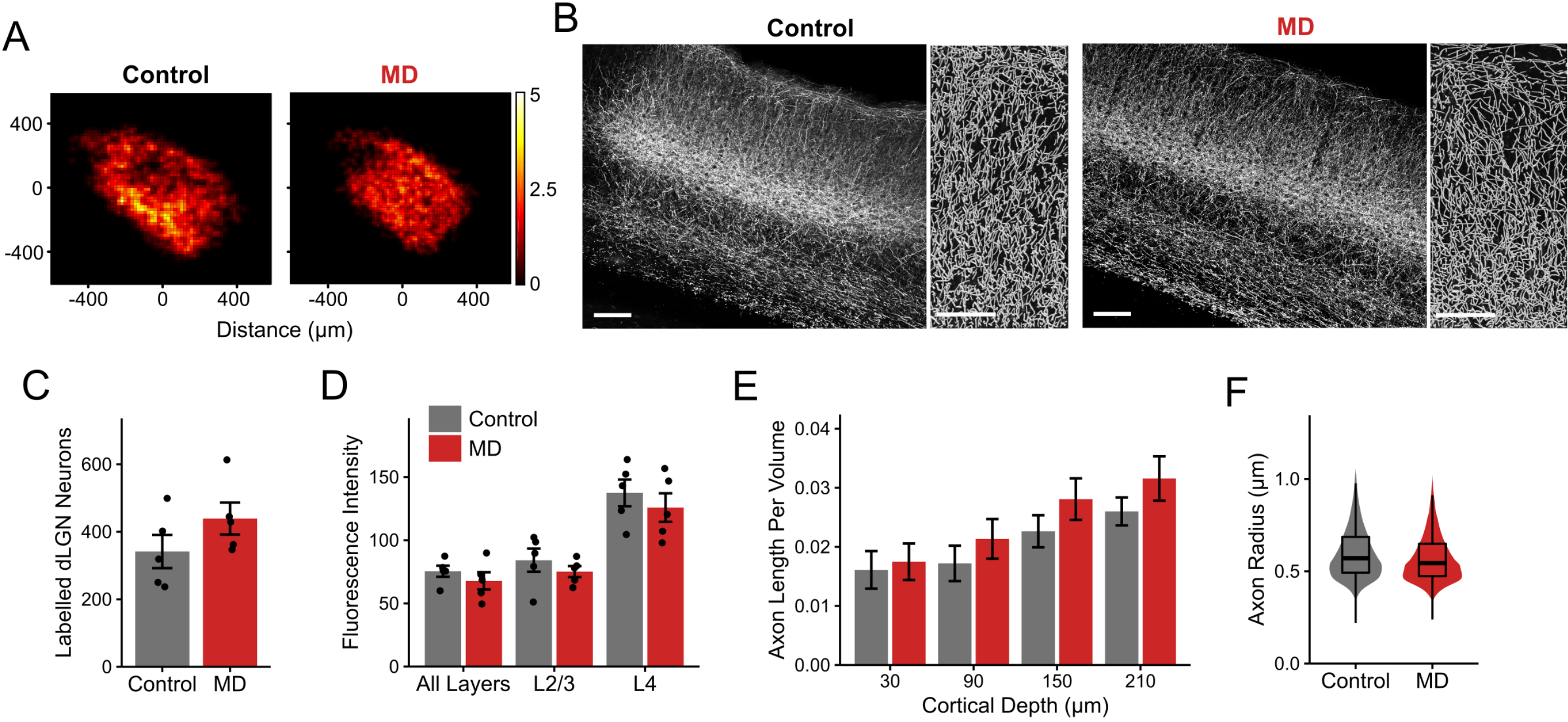
No gross structural loss of thalamocortical connectivity following critical-period MD. **A**, Composite heatmaps showing the spatial distribution of dLGN neurons labeled following GCaMP6s virus injection in control and MD mice. Of ten mice included in this dataset, six were part of the functional data obtained using *in vivo* two-photon calcium imaging. Heatmaps are based on summed cell counts across all sections and mice. **B**, Left: Example fluorescence images (max projection of confocal z-stacks) of V1 coronal sections showing thalamocortical axon labeling in control *vs.* MD mice (scale bar 100 μm). m). Sections were immunostained for GFP. Right: Axons in L1-2/3 were automatically traced, supplemented by manual tracing by a blinded experimenter (scale bar 50 μm). m). Note dense thalamocortical axon labeling in control and MD sections. Example V1 sections and tracing shown are from the same mice shown in Fig. S2A. **C**, Number of dLGN neurons labeled were similar between control and MD mice (mean ± SEM, by animal; Wilcoxon rank sum test: *P* = 0.21). **D**, Mean fluorescence intensity of labeling in V1 sections from control *vs.* MD mice, shown separately for all layers, L2/3 only and L4 only (mean ± SEM, by animal). Mean fluorescence was not significantly different between control and MD mice (2-way ANOVA: effect of MD: P = 0.18, effect of layer: 2.15 × 10^−7^). **E**, Traced axon length per volume (μm). m per μm). m^3^) across different cortical depths in V1 L1-2/3 in control *vs.* MD (mean ± SEM, by section; linear mixed-effects model: effect of MD: *P* = 0.84, effect of cortical depth: *P* = 8.1 × 10^−13^). **F**, Violin plots showing distribution of traced axon radius in V1 L1-2/3 in control *vs.* MD mice. Linear mixed-effects model: control *vs.* MD: *P* = 0.57. In box plots, the central mark indicates the median and the bottom and top edges indicate the 25^th^ and 75^th^ percentiles, respectively. All panels: *n* = 5 control and 5 MD mice, 3 sections per animal.

## DISCUSSION

In this study, we examined the role of early visual experience in shaping visual response properties of dLGN inputs to binocular visual cortex. Our findings confirm the existence of a small proportion of dLGN afferents that are responsive to both eyes (21). We further demonstrate that long-term sensory deprivation (14 days of MD) during the critical period for ocular dominance plasticity leads to a persistent loss of binocular dLGN afferents and significant interocular mismatch of visual properties. Long-lasting changes in ocular dominance, spatial acuity and binocular mismatch following MD have previously been observed for V1 neurons (5, 8, 26). Our findings establish, for the first time, that some of these functional changes (e.g., persistent alterations in ocular dominance and binocular mismatch) are already present at the level of single thalamocortical synapses, indicating that the deficits in binocular integration may begin in the thalamus. On the other hand, we found that overall SF processing in the thalamocortical pathway remains intact in deprived mice, despite marked acuity loss in V1 neurons. These findings provide further evidence for the hypothesis that distinct mechanisms are involved in fine-tuning of neural circuits underlying binocular vision *vs.* spatial acuity (26, 37).

Our result that binocular dLGN afferents exhibited much stronger visual responses compared to monocular dLGN inputs is in agreement with a previous finding demonstrating that binocular dLGN cells receive inputs from a larger number of retinal ganglion cells compared to monocular cells (24). It is currently unknown the mechanisms involved in the development of binocularity and interocular matching of SF and orientation tuning properties in dLGN neurons. Previous work has demonstrated that summed postsynaptic thalamic inputs are already slightly matched in preferred orientation before the critical period, suggesting that dLGN inputs may help shape V1 neurons’ binocular orientation matching during development (38). However, previous work left open the question of how single dLGN neurons achieve binocularity and interocular matching. Our surprising finding that the ipsilateral-(non-deprived) eye responses are placed at a greater disadvantage compared to the contralateral-eye responses, when binocular dLGN boutons become mismatched, hints at an interesting possibility that the ipsilateral input may rely on the contralateral input to guide the matching process. This is in line with previous findings showing that the ipsilateral pathway develops later and is more vulnerable to developmental manipulations compared to the contralateral pathway (2, 3, 39–41).

Taken together, our findings demonstrate that binocular integration in the early feedforward pathway from dLGN to V1 requires normal visual experience during the critical period to develop properly. While it is likely that binocular competition plays a role, the exact locus of action, cell types and molecular factors involved in this developmental mechanism remain to be elucidated (42–44). Considering our results, future studies investigating ocular dominance plasticity and binocular matching will need to disambiguate the relative contributions of thalamic *vs.* cortical mechanisms for binocular integration. Finally, it will be important to determine whether binocular integration in the primate dLGN (23) is also vulnerable to manipulations during the critical period of development, in order to assess the significance of these findings in the context of amblyopia.

## ACKNOWLEDGMENTS

The authors would like to thank Karen Bradshaw and Mariyam Habeeb for technical assistance with preliminary experiments. This work was supported by National Institutes of Health DP2 grant (NEI EY024504) to S.P.G., and Canadian Institutes of Health Research Postdoctoral Fellowship and Knights Templar Eye Foundation Grants to C.Y.H.

## AUTHOR CONTRIBUTIONS

C.Y.H. and S.P.G. conceived experiments, C.Y.H., K.A. and K.J.S. performed experiments, C.Y.H., K.A., K.J.S., J.Z. and C.F. analyzed data, C.Y.H., D.G., J.Z. and J.P.P. built custom software, D.X.F.V. generated preliminary data, C.Y.H. and S.P.G. wrote the manuscript.

## MATERIALS AND METHODS

### Animals

For thalamocortical axon imaging, we used wildtype C57BL/6 mice (Strain No. 027, Charles River) and VGLUT2-Cre mice (Vglut2-ires-cre; Stock No. 016963, Jax Labs). VGLUT2-Cre homozygous mice were bred with wildtype mice to produce heterozygous offspring that were used for imaging. For excitatory V1 neuron imaging, a Camk2a-tTa driver line (Stock No. 007004, Jax Labs) was crossed to a line expressing GCaMP6s under the control of the tetracycline-responsive regulatory element (tetO; Stock No. 024742, Jax Labs) to produce CaMK2a-tTA;tetO-GCaMP6s mice (28); the founder line was heterozygous for both transgenes and maintained by breeding with wildtype mice. Mice were weaned at P19 and co-housed with one or more littermate of the same sex until viral injections. All mice were housed in conventional mouse housing conditions, and both male and female mice were used. For all surgeries, body temperature was maintained at ~37.5°C by a feedback-controlled heating pad and eyes were covered with ophthalmic ointment to prevent drying. All protocols and procedures followed the guidelines of the Animal Care and Use Committee at the University of California, Irvine.

### Monocular deprivation

Mice were monocularly deprived (MD) during the critical period for ocular dominance plasticity (P19 - 33) by eyelid closure (27). Under isoflurane anesthesia (2% for induction, 1 - 1.5% for maintenance), the non-deprived eye was covered with ophthalmic ointment and the other eye was kept moist with sterile saline. Eye lashes were trimmed and upper and lower eyelids were sutured closed using two mattress sutures (7-0 silk, Ethicon). Eyes were checked every 2-3 days for proper closure. On the 14^th^ day of MD, the previously closed eye was reopened and carefully checked for any ocular damage under a microscope. If an eye opened prematurely or was found to be damaged, the animal was excluded from the study. Eye health was further monitored for 1 - 2 weeks following eye reopening.

### GCaMP6s virus delivery

For thalamocortical axon imaging, we initially injected AAV1.Syn.GCaMP6s virus into the dorsolateral geniculate nucleus of the thalamus (dLGN) in wildtype C57BL/6 mice but we found that this approach led to labeling of some V1 cell somata. Thus, we employed another approach of injecting AAV1.Syn.Flex.GCaMP6s virus into dLGN in VGLUT2-Cre mice. Since vesicular glutamate transporter 2 (VGLUT2) is predominantly expressed by thalamic neurons (45), we were able to restrict GCaMP6s expression specifically to dLGN neurons using this approach, with little to no cells being labeled in V1. Results from the two approaches were similar and data from 3 wildtype and 8 VGLUT2-Cre mice used for functional imaging were combined for analysis. Viral vectors were obtained from Penn Vector Core.

For dLGN injections, mice (P58 - 80; mean: P67) were placed in a stereotaxic frame under isoflurane anesthesia (2% for induction, 1 - 1.5% for maintenance). Mice were injected with lactated Ringer’s solution and carprofen (5 mg/kg, s.c.) for hydration and analgesia. Scalp was retracted and a small burr hole was made at the injection site using a pneumatic drill. Coordinates used for targeting dLGN was ~2.2 mm posterior, ~2.2 mm lateral from bregma, and ~2.6 mm deep from the brain surface. Viral vectors diluted to the final titre of ~1 × 10^12^ GC/ml were loaded into a glass pipette and injected into dLGN in one hemisphere (total volume: 80 nl, rate: 8 nl/min). In MD mice, the hemisphere contralateral to the deprived eye was injected. The skull and injection site were kept moist with saline during the injection. Following surgery, mice were placed on a heat pad to recover and monitored for post-operative health.

### Cranial window implantation

Headplate attachment and craniotomy were performed in one surgery following previously reported procedures (29). Briefly, mice were anesthetized with isoflurane (2% for induction, 1 - 1.5% for maintenance) and topical lidocaine (2%) was applied to provide analgesia. With the head secured in a stereotaxic frame, the skull was exposed and an approximate location of bV1 was marked. A layer of cyanoacrylic glue (3M Vetbond™) was applied to the skull and a custom-printed black headplate was centered over bV1 and fixed to the skull using black dental acrylic (Ortho-Jet, Lang Dental) at an angle parallel to the imaging site. A craniotomy (4 or 5 mm dia.) was performed and a No. 1 glass coverslip was placed over the exposed brain and sealed with cyanoacrylic glue and dental acrylic. Mice were placed in a warm cage to recover until mobile and given daily injections of lactated Ringer’s and carprofen for at least 3 days and monitored for post-operative health. In MD mice, craniotomy was performed over bV1 contralateral to the deprived eye. Age of mice at craniotomy was P83 - 106 (mean: P94).

### Widefield imaging for bV1 mapping

Widefield imaging for bV1 mapping was performed through the cranial window after ≥ 4 days of recovery following craniotomy and age of mice at mapping was P90 - 114 (mean age: P101). For mice used for thalamocortical axon imaging, mapping of binocular V1 was performed using widefield intrinsic signal imaging, following published procedures (27, 29). Briefly, awake mice were placed on a smooth platform, head-fixed and shown contrast-reversing noise stimulus that spanned central 30° of the mouse’s visual field. The stimulus was swept either up or down periodically every 20 s. The stimulus was generated by multiplying a band-limited (<0.05 cpd, >2 Hz) binarized spatiotemporal noise movie with a one-dimensional Gaussian spatial mask (30°) using custom Python scripts. Visual stimuli were presented on a gamma-corrected 24” LED monitor (ASUS VG248, 60 Hz refresh rate, 20 cd/m^2^ mean luminance) at a viewing distance of 25 cm. Widefield fluorescence images were acquired using a SciMedia THT macroscope (Leica PlanApo 1.0X, 6.5 × 6.5 mm imaging area) equipped with an Andor Zyla sCMOS camera. For visualizing vasculature, a green (530 nm) LED was used. The camera was focused ~600 μm). m beneath the brain surface, located using vasculature, and intrinsic signals were acquired with a red (617 nm) LED. The stimulus was presented for 5 min under binocular viewing conditions and typically 2 - 3 repeats were run for each condition. Data were analyzed to extract maps of amplitude and phase of cortical responses by Fourier analysis at the frequency of stimulus repetition (46) using custom MATLAB (MathWorks) software. Amplitude was computed by taking the maximum of the Fourier amplitude map smoothed with a 5x5 Gaussian kernel. For Cam2k-tTA;tetO-GCaMP6s transgenic mice, mapping of bV1 was performed using widefield GCaMP6s imaging (blue LED excitation at 465 nm), following procedures published previously (29).

### *In vivo* two-photon Ca^2+^ imaging

All imaging was performed in awake head-fixed mice sitting on a smooth tablet surface. Mice were habituated on the imaging setup for 0.5 - 1 hour each day for 1 - 2 days prior to imaging. From the same mouse, imaging was performed typically for 2 - 3 hours per day for 2 - 5 days from different fields of view (age at imaging: P93 - 132, mean: P106). The average time interval between GCaMP6s virus injection and two-photon imaging was 39 days.

A resonant two-photon microscope (Neurolabware) and 920 nm excitation laser (Mai Tai HP, Spectra-Physics) were used for GCaMP6s imaging, following previously published procedures (29) with modifications. A Nikon 16X (NA = 0.8) water-immersion objective was used. For dLGN bouton imaging, fields of view typically covered ~220 μm × 260 μm). m and image sequences were acquired at 8 Hz (990 lines) at depths of 140 ± 37 μm). m (mean ± SD in 34 fields) below the pia, corresponding to cortical layers 1−2/3. Recordings were confined to anterior and middle parts of bV1. For V1 excitatory neuron recordings, fields were typically ~700 μm). m × 500 μm). m, acquired at 7.7 Hz (1024 lines) and recordings were performed in middle bV1 at cortical depths of approximately 200 μm). m, corresponding to L2/3. Data acquisition was controlled by Scanbox software (Scanbox).

Visual stimuli were generated by custom Python software using PsychoPy 1.8 library. Spherically corrected stimuli were presented on a gamma-corrected 24” LED monitor (Asus VG248, 60 Hz refresh rate, 20 cd/m^2^), placed at 25 cm from the mouse’s eyes. The stimuli included full-field drifting sinusoidal gratings (contrast: 99%) of 5 - 6 spatial frequencies (SF; 0.03 - 0.48 or 0.03 - 0.96 cpd, spaced logarithmically) and 8 directions (0 - 315°, in 45° steps) at a temporal frequency of 2 Hz, a blank (uniform luminance) condition, and a full-field flicker (2 Hz) condition. Each trial consisted of a visual stimulus for 2 s and a uniform gray screen for 2 s. Different stimuli were presented in a random order without replacement and typically 8 repeats were run per stimulus condition. Visual stimuli were presented to one eye at a time, either first to the contralateral or ipsilateral eye using an occluder, and the order of eye presentation was chosen randomly for each session. Eyes were monitored using IR-compatible GigE cameras (Allied Vision Mako-131B). The illumination by the infrared laser (used for two-photon imaging) was used for pupil tracking.

### Ca^2+^ imaging data analysis

Custom Python software was used to remove motion artifacts, manually identify dLGN boutons and cells, extract fluorescence traces, and perform batch analyses, according to previously described procedures (29) with modifications. We implemented a motion correction algorithm that corrects for translational artifacts by minimizing the Euclidean distance between frames and a template image, using a Fourier transform approach (47). The outcome of the motion correction was checked by visualizing the mean intensity of 40 pixels in the middle of the frame throughout the movie. To identify regions of interest (ROIs) as boutons or cell bodies, we used the summed intensity projection of the motion-corrected movies and applied morphological criteria to manually identify them.

All pixel values within the ROI region were averaged to yield the fluorescence trace for the ROI. The fluorescence signal of a cell body at time *t* was determined (48, 49) as follows: *F*_*cell*_ (*t*)=*F*_*soma*_ (*t*) −(*R*· *F*_*neuropil*_ (*t*)). R was empirically determined to be 0.7 by comparing blood-vessel intensity of GCaMP6s signal to that in the neuropil. The neuropil signal was estimated by taking the mean of the signal in all pixels within ~3 μm). m radius outside the cell’s outline. Bouton data were treated to a similar neuropil subtraction except that for neuropil, a radius of ~1 μm). m outside the bouton’s outline was used.

To determine a ROI’s response to each stimulus trial, the ROI’s trace during the stimulation period was first normalized to the baseline fluorescence value averaged over the 0.5 s preceding the stimulus (ΔF / F_0_). Then, the mean response amplitude (mean ΔF / F_0_) was generated for each stimulus type by averaging the normalized response across all trials of that stimulus. A ROI’s spontaneous calcium fluctuation was estimated using the ROI’s mean response amplitude during blank stimulus presentation. For each SF, a ROI’s responsiveness was determined using a one-way ANOVA (*P* < 0.01) across responses for all orientations for that SF against responses for the blank condition. For most of the analyses in this paper, we restricted our analyses to ROIs whose responses at the peak SF (SF that gave the strongest response) reached statistical significance at *P* < 0.01 (except for data depicted in Fig. 1F ; see below for ODI calculation). In Fig. S1F-G, we explored whether lowering or raising the significance level to *P* < 0.05 or *P* < 0.005 affected our results and we found that the effect of MD on binocular bouton number and proportion remained statistically significant under the different criteria. For V1 L2/3 neuron recordings, an additional criterion was placed such that only cells whose mean ΔF / F_0_ for their preferred stimulus (R_pref_) was ≥ 0.05 were included for further analyses.

For each ROI, the preferred orientation (θ_pref_) was determined at the ROI’s peak SF, by calculating half the mean of the directional vectors weighted by the response F(θ) at each orientation as follows:

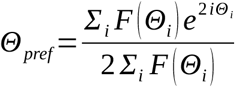

For each SF, an orientation tuning curve was obtained by fitting a sum of Gaussians function on mean response amplitudes for the eight orientations. The response amplitude at the preferred orientation based on the fitted values was designated as R(θ_pref_). To fit a SF tuning curve, response amplitudes at the preferred orientation (θ_pref_) across the spatial frequencies were fitted with a Gaussian function. The bandwidth was calculated by taking the square root of the width at half the maximum of the fit. R_pref_ is the mean amplitude of the ROI’s response to its preferred grating (preferred orientation and SF).

Orientation and direction selectivity for a ROI was determined using a method based on circular variance of the cell’s response as follows:

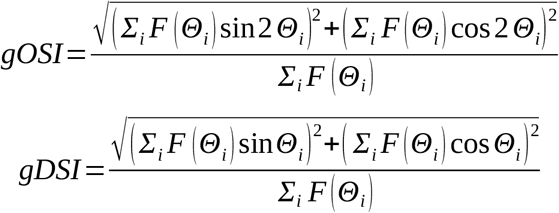

Ocular dominance index (ODI) for each ROI was calculated as (C – I) / (C + I), where C is R_pref_ for contralateral-eye responses and I is R_pref_ for ipsilateral-eye responses. In cases where no significant response was detected for one of the eyes according to the responsiveness criteria described above, R_pref_ for that eye was set to zero. Thus, responses that were purely driven by the contralateral-*vs.* ipsilateral-eye stimulation were given ODI values of 1 and -1, respectively. The method of estimating ODI differed for data depicted in Fig. 1F only; color-coding was based on ODI values calculated according to the same formula as above, except that if one eye did not meet the responsiveness criteria, its R_pref_ was not set to zero. Thus, if one of the eyes’ responses passed the responsiveness criteria, the other eye’s R_pref_ was used to calculate ODI in Fig. 1F.

### Histological procedures and anatomical data analysis

After the last imaging session, mice were anesthetized and transcardially perfused with saline and 4% paraformaldehyde. Age of mice at perfusion was P112 - 142 (mean: P119). Brains were extracted, post fixed and cryoprotected with 30% sucrose. The brain was sectioned coronally in 50 μm). m using a frozen sliding microtome (Microm HM450, Thermo Scientific). Tissue was processed for GFP immunostaining in free floating sections as follows. Sections were blocked for 1 hour at room temperature with 0.5% Triton-X (T8787, Sigma) and 10% BSA (BP1600-100, Fisher) in PBS, then incubated overnight at room temperature with chicken anti-GFP antibody at 1:500 dilution (GFP-1020, Aves Labs). Sections were then washed in PBS and incubated for 2 hours at room temperature with goat anti-chicken IgG antibody tagged with Alexa-488 at 1:1000 dilution (A-11039, Life Technolgies). Sections were further processed for nuclear staining (Hoechst 33342), washed in PBS, coverslipped with Flouromount-G (Southern Biotech) and imaged.

For dLGN sections, we used an epifluorescence microscope (Zeiss Axio Imager 2) with a 10X objective. For cell counting, labeled cells in dLGN sections every 200 μm). m (3 sections per animal) were manually counted using the cell counter plugin in Fiji. Total number of labeled dLGN neurons as well as the spatial distribution of labeled neurons in dLGN was quantified for each animal. Functionally imaged brains where post-hoc anatomical data revealed that cells were labeled in the neighboring thalamic nucleus LP were excluded from analysis.

For V1 sections, we first took images using the epifluorescence microscope with a 10X objective. Cortical layers were identified using nuclear staining. In order to estimate thalamocortical axon density, we obtained the mean fluorescence intensity across the cortical depths in a densely labeled area of a fixed size in V1 (186 μm). m horizontal × 932 μm). m vertical) of each section and quantified labeling intensity in each layer (Fig. 4D). To get a more accurate estimate of the axon density, we sought to segment axons from the images. For this, z-stack images were taken of V1 sections every 200 μm). m (3 sections per animal) using a Zeiss LSM700 confocal microscope and a 20X objective (NA = 1.0). Images were rotated and cropped to include only superficial layers (L1 and L2/3) in a densely labeled volume of a fixed size in V1 (100 μm). m horizontal × 242 μm). m vertical × all z slices), which corresponds to the layers that were functionally imaged using *in vivo* two-photon Ca^2+^ imaging. Open source neuron tracing software neuTube (50) with custom modifications was used to detect axons automatically. The output traces were filtered to remove abnormally large radius nodes, branching points and isolated nodes. From visual inspection, the auto-segmentation did not detect all visible axons, so the tracing was supplemented by manual tracing by a blinded experimenter. From this final set of traces, we quantified the total axon length per volume and axon radius of thalamocortical axons in V1 L1-2/3 (3 sections per animal).

### Statistical analysis

The statistical determination of visual responsiveness is described in detail above; the ANOVA tests for responsiveness, curve-fitting for orientation and SF tuning and related selectivity/bandwidth calculations were performed in custom Python routines. Plotting of fluorescence traces, example tuning curves and overlay of vector graphics on images were done using MATLAB or Python scripts. All other statistical analyses and data plotting were performed using custom software in R. In addition to conventional statistics (Chi-squared tests, T-tests, Wilcoxon rank sum tests, 2- and 3-way ANOVAs, Kolmogorov-Smirnov tests), multilevel statistics were employed in some cases to take into account the nested design of our data (*e.g.*, boutons, neurons, sections nested inside mice). Multilevel linear mixed-effects models with Satterthwaite’s approximation were used, with experimental variables (*e.g.*, control *vs.* MD) as fixed variables and mouse ID as a random variable. For each analysis, the exact statistical test used and sample sizes are described in figure legends. All tests were two-tailed. Data are reported as mean ± SEM unless otherwise noted.

