## Supplemental Information for "Critical-Period Visual Deprivation Disrupts Binocular Integration but Spares Spatial Acuity in the Geniculocortical Pathway"

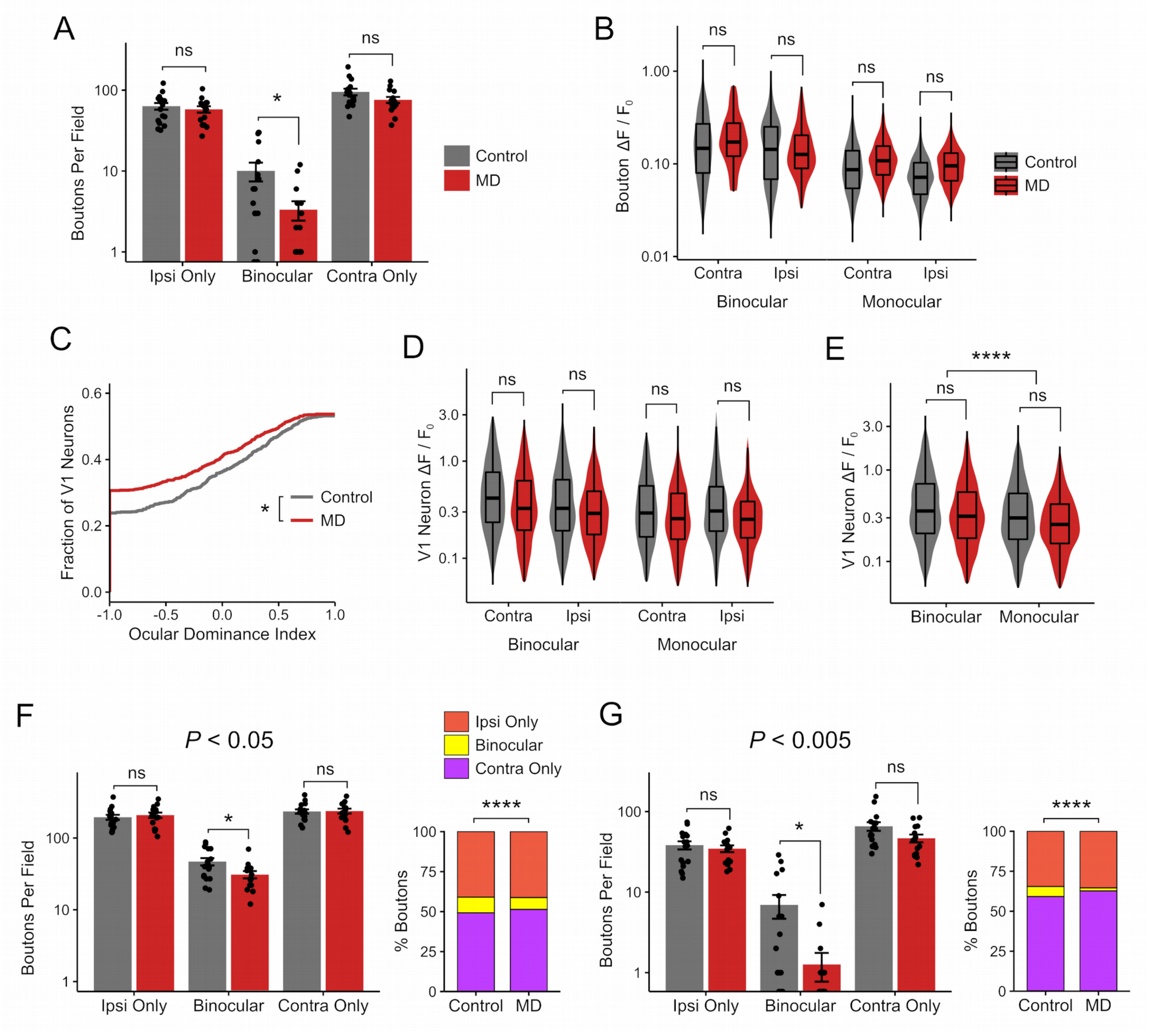
A
Boutons Per Field
ns
*
ns
Control
MD
Ipsi Only
Binocular
Contra Only
B
Bouton ΔF / F0
Contra
Ipsi
Binocular
Monocular
C
Fraction of V1 Neurons
Ocular Dominance Index
D
V1 Neuron ΔF / F0
E
****
F
P < 0.05
Ipsi Only
Binocular
Contra Only
% Boutons
G
P < 0.005

**Figure S1**

**Figure S1. Loss of binocularity in dLGN boutons and V1 neurons following critical-period MD (corresponds to Figure 1).**

(A) Number of visually responsive boutons per field in control *vs.* MD mice (mean ± SEM per field, $n$ = 17 control, 15 MD fields; linear mixed-effects model, effect of MD for ipsi-only: $P$ = 0.62, binocular: $P$ = 0.02, contra-only: $P$ = 0.08). (B) Violin and overlaid box plots of mean response amplitude of boutons (linear mixed-effects model, effect of MD: $P$ = 0.49, binocular *vs.* monocular: $P = 2.2 \times 10^{-16}$, contra *vs.* ipsi: $P = 2.2 \times 10^{-11}$). (C) Cumulative distribution of ODI values from V1 L2/3 neurons, showing a shift towards the ipsilateral eye in MD mice (ODI: -1 means ipsi-only, +1 means contra-only; Kolmogorov-Smirnov test, $P$ = 0.01). (D) Violin and overlaid box plots of mean response amplitude of V1 neurons (linear mixed-effects model, effect of MD: $P$ = 0.67, binocular *vs.* monocular: $P = 3.1 \times 10^{-13}$, contra *vs.* ipsi: $P$ = 0.06). (E) Violin and overlaid box plots of mean response amplitude of V1 neurons, grouped into binocular *vs.* monocular (linear mixed-effects model, effect of MD: $P$ = 0.71, binocular *vs.* monocular: $P = 2.75 \times 10^{-14}$). Note that similar to dLGN boutons (Fig. 1J and Fig. S1B), binocular V1 neurons showed larger response amplitudes compared to monocular neurons. (F-G) The binocular bouton loss following critical-period MD (corresponds to Fig. 1M and Fig. S1A) is shown using two additional statistical criteria in determining visual responsiveness: more liberal (F: $P < 0.05$) or more conservative (G: $P < 0.005$) criteria than the typical criterion used in this study ($P < 0.01$; see STAR Methods). Linear mixed-effects model: effect of MD for ipsi-only: $P$ = 0.74 (F), $P$ = 0.62 (G), for binocular: $P$ = 0.01 (F), $P$ = 0.02 (G), for contra-only: $P$ = 0.99 (F), $P$ = 0.05 (G). Chi-squared test: for F, $\chi^2(2) = 33.7$, $P = 4.7 \times 10^{-8}$, $n$ = 8099 *vs.* 9307 visually responsive boutons in total, 10% *vs.* 7% were binocular (control *vs.* MD); for G, $\chi^2(2) = 45.5$, $P = 1.3 \times 10^{-10}$, $n$ = 1887 *vs.* 1881 visually responsive boutons in total, 6% *vs.* 2% were binocular (control *vs.* MD). Note that the effect of binocular bouton loss in MD mice remained statistically significant under different data inclusion criteria. Sample sizes for A-E: Boutons: $n$ = 2866 boutons in 5 control mice, 2975 boutons in 6 MD mice; V1 neurons: $n$ = 1051 neurons in 9 control mice, 1145 neurons in 4 MD mice. In box plots, the central mark indicates the median and the bottom and top edges indicate the 25$^{th}$ and 75$^{th}$ percentiles, respectively. All panels: $^{ns}P > 0.05$, $^{*}P < 0.05$, $^{****}P < 0.0001$.

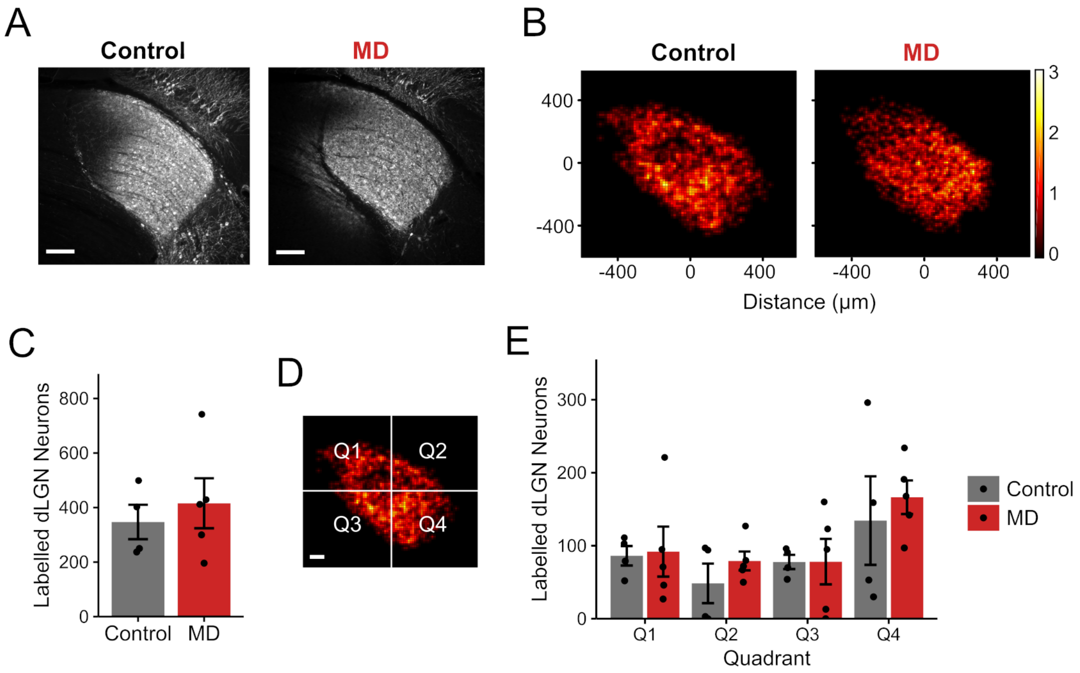


**Figure S2. GCaMP6s-labelling in dLGN is comparable between control and MD mice used for functional thalamocortical axon imaging (corresponds to Figure 1).**
(A) Example fluorescence sections of dLGN neurons labeled following GCaMP6s virus injection in control and MD mice that were used for *in vivo* two-photon $Ca^{2+}$ thalamocortical axon imaging. Sections were immunostained for GFP (scale bar 100 μm). Example dLGN sections shown are from the same mice shown in Fig. 4B. (B) Composite heatmaps showing the spatial distribution of labeled dLGN neurons in control and MD mice. Heatmaps are based on summed cell counts across all sections and mice. (C) Numbers of dLGN neurons labeled were similar between functionally imaged control *vs.* MD mice (mean ± SEM of by-animal values; T-test: $P = 0.56$). (D) Quadrants used in quantifying the spatial distribution of labeled dLGN neurons (scale bar 100 μm). (E) Numbers of labeled dLGN neurons were similar between control and MD mice across all quadrants (mean ± SEM of by-animal values, 2-way ANOVA: control *vs.* MD: $P = 0.56$, effect of quadrant: $P = 0.03$, interaction: $P = 0.93$). All panels: $n = 4$ control and 5 MD mice, cells counted on three sections per animal.

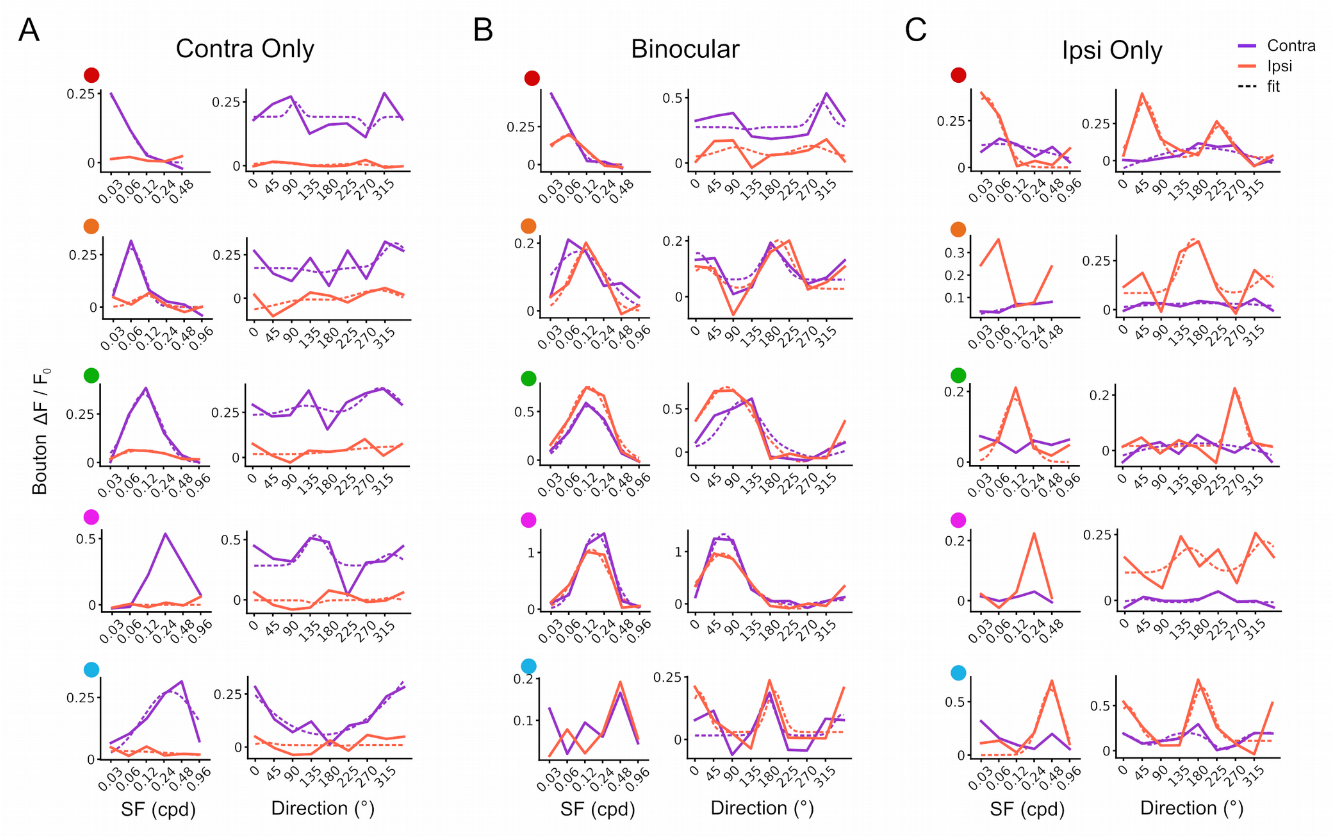


**Figure S3. Example SF and orientation tuning curves of dLGN boutons in control mice (corresponds to Figures 2 and 3).**

(A-C) Each pair of plots shows spatial frequency (SF) tuning curve (left) and orientation tuning curve at peak SF (right) of the same bouton. Five example boutons for each group (A: contra-only monocular, B: binocular, C: ipsi-only monocular) are shown. Examples are ordered such that boutons with the lowest preferred SF are placed at the top and those with the highest preferred SF are shown at the bottom. For all tuning curves, mean response amplitude ($\Delta F/F_0$) is plotted on the Y axis. Purple traces are from contralateral-eye trials; orange traces are from ipsilateral-eye trials; solid traces are mean response amplitudes; dotted traces are fitted curves based on the mean values. Fits are omitted if curve-fitting failed to converge. Dots to the left are color-coded according to peak SF of the bouton (red: 0.03 cpd, orange: 0.06 cpd, green: 0.12 cpd, magenta: 0.24 cpd, light blue: 0.48 cpd; peak SF of the contralateral-eye response was used for color-coding of binocular boutons).

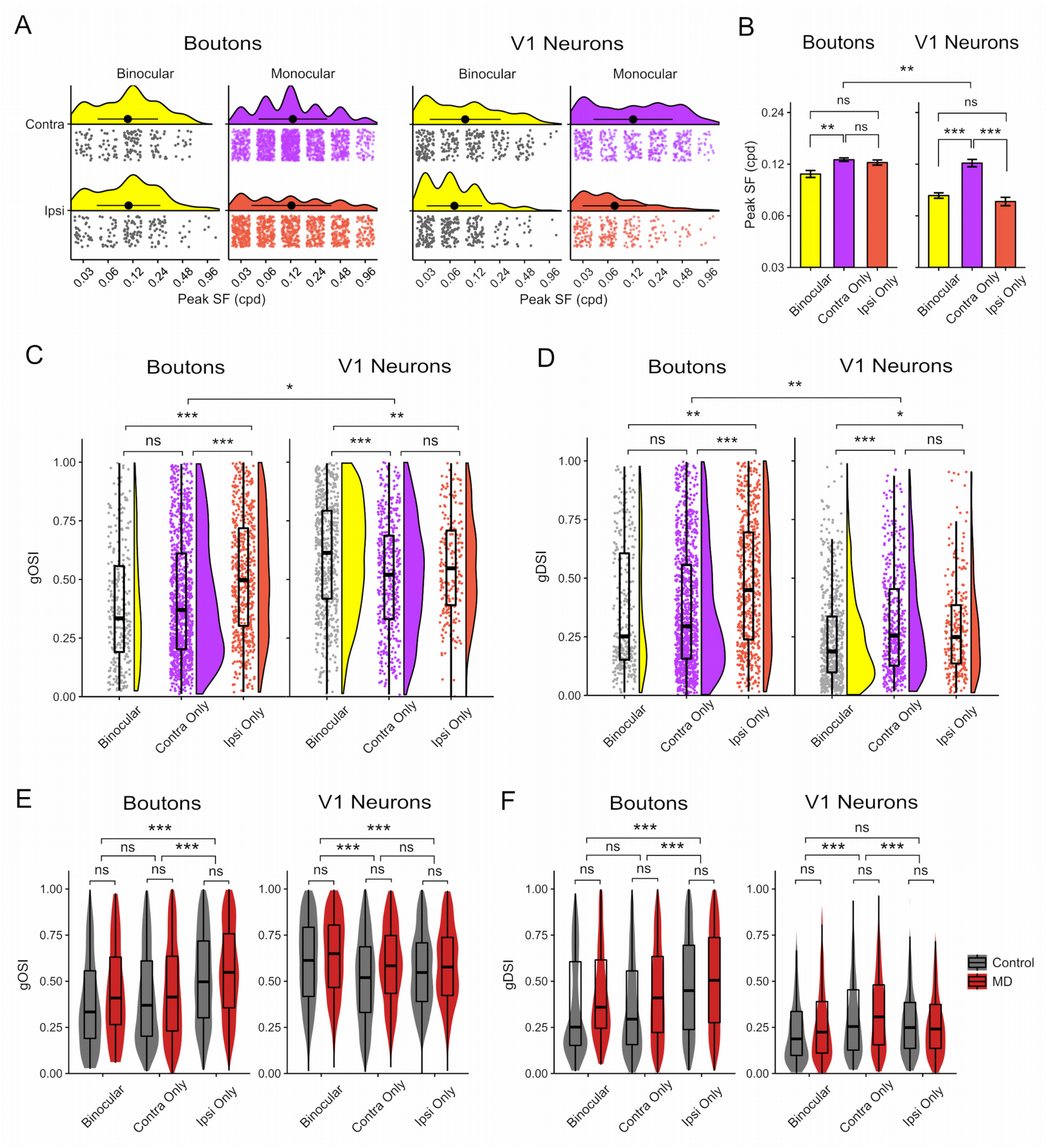

A
Boutons
V1 Neurons
Binocular
Monocular
Contra
Ipsi
Peak SF (cpd)
B
Peak SF (cpd)
Binocular
Contra Only
Ipsi Only
C
gOSI
D
gDSI
E
F
Control
MD


**Figure S4**

**Figure S4. Comparison of visual properties of dLGN boutons *vs.* V1 neurons (corresponds to Figures 2 and 3).**

(A-D) Control data only. (A) Rain cloud plots showing distributions of peak SF in contra-only monocular (purple), ipsi-only monocular (orange) and binocular (yellow with gray dots) dLGN boutons (left) and V1 neurons (right). Black filled circles and lines indicate mean ± SD. (B) Bar plots (error bars: mean ± SEM) of peak SF in binocular, contra-only *vs.* ipsi-only monocular dLGN boutons (left) and V1 neurons (right). Boutons: linear mixed-effects model, binocular *vs.* contra-only: $P = 0.004$, binocular *vs.* ipsi-only: $P = 0.15$, contra-only *vs.* ipsi-only: $P = 0.12$. V1 neurons: linear mixed-effects model, binocular *vs.* contra-only: $P = 9.7 \times 10^{-11}$, binocular *vs.* ipsi-only: $P = 0.21$, contra-only *vs.* ipsi-only: $P = 9.8 \times 10^{-10}$, dLGN boutons *vs.* V1 neurons: $P = 0.006$. (C) Rain cloud plots showing distribution of gOSI values in binocular, contra-only *vs.* ipsi-only monocular boutons (left) and V1 neurons (right). Boutons: linear mixed-effects model, binocular *vs.* contra-only: $P = 0.08$, binocular *vs.* ipsi-only: $P = 7.5 \times 10^{-11}$, contra-only *vs.* ipsi-only: $P = 9.7 \times 10^{-5}$. V1 neurons: linear mixed-effects model, binocular *vs.* contra-only: $P = 1.1 \times 10^{-6}$, binocular *vs.* ipsi-only: $P = 0.001$, contra-only *vs.* ipsi-only: $P = 0.23$, dLGN boutons *vs.* V1 neurons: $P = 0.01$. (D) Rain cloud plots showing distribution of gDSI values in binocular, contra-only *vs.* ipsi-only monocular boutons (left) and V1 neurons (right). Boutons: linear mixed-effects model, binocular *vs.* contra-only: $P = 0.72$, binocular *vs.* ipsi-only: $P = 0.004$, contra-only *vs.* ipsi-only: $P = 5.2 \times 10^{-9}$. V1 neurons: linear mixed-effects model, binocular *vs.* contra-only: $P = 1.4 \times 10^{-6}$, binocular *vs.* ipsi-only: $P = 0.01$, contra-only *vs.* ipsi-only: $P = 0.25$, dLGN boutons *vs.* V1 neurons: $P = 0.005$. (E) Violin and overlaid box plots of gOSI in binocular, contra-only *vs.* ipsi-only monocular dLGN boutons (left) and V1 neurons (right) in control and MD mice. Boutons: linear mixed-effects model, effect of MD: $P = 0.42$, binocular *vs.* contra-only: $P = 0.58$, binocular *vs.* ipsi-only: $P = 4.47 \times 10^{-10}$, contra-only *vs.* ipsi-only: $P = 4.27 \times 10^{-11}$. V1 neurons: linear mixed-effects model, effect of MD: $P = 0.31$, binocular *vs.* contra-only: $P = 1.43 \times 10^{-8}$, binocular *vs.* ipsi-only: $P = 3.58 \times 10^{-5}$, contra-only *vs.* ipsi-only: $P = 0.35$. (F) Violin and overlaid box plots of gDSI in binocular, contra-only *vs.* ipsi-only monocular dLGN boutons (left) and V1 neurons (right) in control and MD mice. Boutons: linear mixed-effects model, effect of MD: $P = 0.32$, binocular *vs.* contra-only: $P = 0.91$, binocular *vs.* ipsi-only: $P = 0.0003$, contra-only *vs.* ipsi-only: $P = 7.63 \times 10^{-12}$. V1 neurons: linear mixed-effects model, effect of MD: $P = 0.36$, binocular *vs.* contra-only: $P = 8.9 \times 10^{-11}$, binocular *vs.* ipsi-only: $P = 0.17$, contra-only *vs.* ipsi-only: $P = 0.0001$. In box plots, the central mark indicates the median and the bottom and top edges indicate the 25$^{th}$ and 75$^{th}$ percentiles, respectively. Sample sizes: $n = 2866$ boutons in 5 control mice, 2975 boutons in 6 MD mice; 1051 neurons in 9 control mice, 1145 neurons in 4 MD mice. All panels: $^{ns}P > 0.05$, $^{*}P < 0.05$, $^{**}P < 0.01$, $^{***}P < 0.001$.

**Supplemental Movie 1. *In vivo* two-photon $Ca^{2+}$ imaging of dLGN axons in V1 (corresponds to Figure 1).**

The movie shows visually evoked activity of V1 projecting axons from dLGN neurons, visualized using GCaMP6s. The recording shown is from L1-2/3 of binocular V1 in a control mouse. During the recording, the mouse was awake, viewing a series of drifting gratings with various spatial frequencies and orientations. The field of view corresponds to the example field shown in Fig. 1E-F. Note that the axon coursing through the center of the field displayed significant visual responses to both eyes (*i.e.*, binocular).
